# ABC Transporters are billion-year-old Maxwell Demons

**DOI:** 10.1101/2021.12.03.471046

**Authors:** Solange Flatt, Daniel M. Busiello, Stefano Zamuner, Paolo De Los Rios

## Abstract

ABC transporters are a broad family of biological machines, found in most prokaryotic and eukaryotic cells, performing the crucial import or export of substrates through both plasma and organellar membranes, and maintaining a steady concentration gradient driven by ATP hydrolysis. Building upon the present biophysical and biochemical characterization of ABC transporters, we propose here a model whose solution reveals that these machines are an exact molecular realization of the Maxwell Demon, a century-old abstract device that uses an energy source to drive systems away from thermodynamic equilibrium. In particular, the Maxwell Demon does not perform any direct mechanical work on the system, but simply selects which spontaneous processes to allow and which ones to forbid based on information that it collects and processes. In the molecular model introduced here, the different information-processing steps that characterize Maxwell Demons (measurement, feedback and resetting) are features that emerge from the biochemical and structural properties of ABC transporters, allowing us to develop an explicit bridge between the molecular level description and the higher-level language of information theory.

## Introduction

Transport processes across membranes are crucial in every living organism^1^: the ability to import or export substrates is essential for example to absorb nutrients or to expel metabolic waste, toxins or drugs. ATP-Binding Cassette (ABC) transporters represent a very broad family of transporters that are found in most prokaryotic and eukaryotic cells^2, 3^. They are transmembrane proteins whose main function is the active (*i*.*e*. ATP-hydrolysis driven) import or export of selected substrates both through the plasma and through organellar membranes (*e*.*g*. between cytosol and *endoplasmic reticulum*). The major distinction in the ABC family is between importers, that allow the intake of molecules from the environment into the cell, and exporters, which transport molecules in the reverse direction. The dependence of ABC transporters on ATP as an energy source does not come as a surprise, because transport often takes place against concentration gradients, thus against equilibrium thermodynamics, and energy must be invested to support a steady substrate concentration difference across the membrane. Overall, the functional cycle of ABC transporters requires them to be in a conformation suitable to bind substrates on one side of the membrane, then switching conformation upon substrate binding that stimulates ATP hydrolysis, and, after substrate release and nucleotide exchange, switching back to the previous conformation, ready to start a new transport cycle (Fig. 1A)^3^. Although ABC transporters have been extensively characterized both structurally and biochemically^4^, a comprehensive framework that integrates the available information into a simple, instructive and physically consistent model is still lacking.

**Figure 1.**
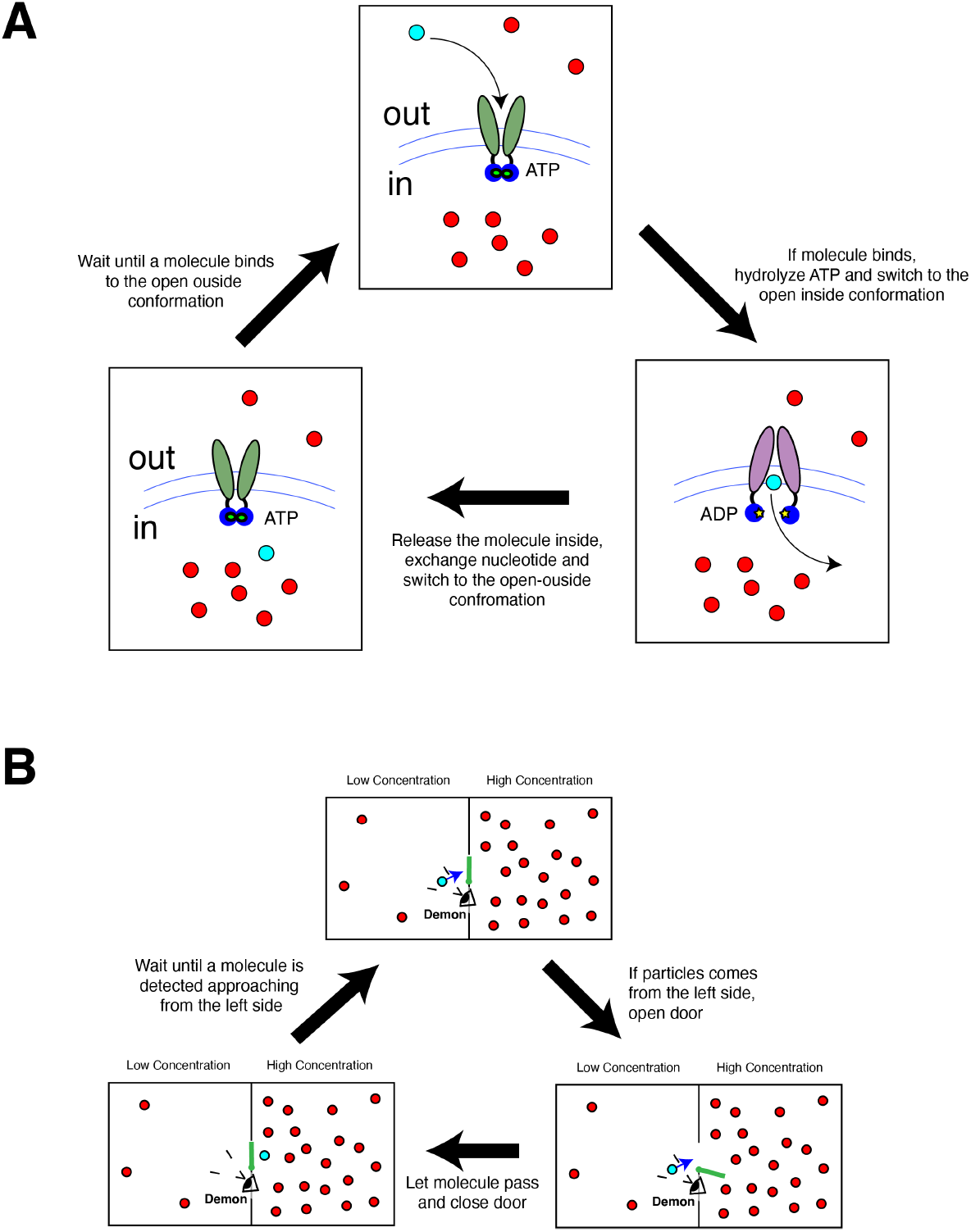
ABC Transporters and Maxwell Demons. A) In a simplified description, ABC transporters (here an importer) switch from an open-outside conformation bound to ATP to an open-inside conformation upon substrate binding and subsequent ATP hydrolysis. After substrate release and nucleotide exchange, the transporter switches back to the open-outside conformation ready for a new cycle (for a more detailed model, see Fig.2). B) When the Maxwell Demon detects a molecule approaching the door from the left, it opens the door and let it pass, closing the door afterwards, and waiting for a new particle to approach.

Because of their ability to maintain a concentration difference between the two sides of a membrane, ABC transporters are reminiscent of *Maxwell Demons*^5^. The Maxwell Demon is an idealized agent (device) that was proposed originally as a challenge to the second law of thermodynamics. In a nutshell, a box is divided in two halves separated by a wall with a door, and is filled with molecules (see Fig. 1B). If the door is open, the molecules will eventually reach their equilibrium state and distribute evenly in the two halves. The Demon operates (opens or closes) the door, and it does so depending on the presence of approaching molecules. If it detects a molecule coming from, say, the left side, it opens the door and let it pass. In the absence of molecules, or if molecules arrive to the door from the right, the Demon will keep the door shut. Repeating this action over and over again, the final result is, intuitively, an accumulation of molecules in the right half of the box, thus establishing a concentration gradient. Since a steady concentration gradient is at odds with thermodynamic equilibrium, it can be sustained only by the constant investment of energy by the Demon. Yet, the Demon does not perform any direct work on the molecules, for example by forcefully dragging them from one side to the other of the door. It only selects which random door crossings, driven solely by thermal motion, are allowed, and which ones are not (this is also sometimes called a *Brownian ratchet*^6^). The solution to this paradox, namely the persistence of a steady non-equilibrium setting without any apparent energy consumption, lies in realising that the Maxwell Demon is actually an information-processing device going through three basic steps^7^. First it must collect information (*measurement*: is there a particle coming from the correct side?). The result of the measurement triggers the *feedback*, which consists in writing the measurement outcome in a physical memory (say, one bit: 0=molecule absent, 1=molecule present), to be then used to decide the state of the door (0=keep the door shut, 1=open the door); eventually, the memory and the door must be reset to allow the Demon to go through a new cycle (*resetting*: close the door and set the bit to 0). This causal, directed sequence of information-processing steps requires energy because it would be otherwise incompatible with equilibrium reversibility.

In this work we present a model of ABC transporters that is based *solely* on present available structural and biochemical information. Its solution reveals that the biochemical conditions that allow ABC transporters to sustain a concentration gradient against equilibrium exactly match the information-processing steps highlighted above. Thus, ABC transporters *are* Maxwell Demons that emerged from billions of years of molecular evolution.

This formal correspondence further allowed us to quantify the information stored and erased during one full cycle of Demon operations (measurement-feedback-resetting), showing that the total processed information is, in fact, intimately connected to the steady concentration gradient across the membrane. This approach highlights the role of information processing by ABC transporters, and from a more general perspective, reveals that the high-level information-theory description of biological processes is an emergent property of their detailed non-equilibrium molecular description.

## Results

### A structurally and biochemically based model of ABC transporters

Structurally, ABC Transporters are dimers constituted by two (usually) identical monomers, each comprising two main subunits: a Transmembrane Domain (TMD) and a Nucleotide Binding Domain (NBD)^4, 8^. The TMDs (two in the dimer) span the membrane and create the channel for the translocation of substrates^3^. The NBDs are located inside the cell, and constitute the binding site of the nucleotides, either ATP or ADP. For the sake of simplicity, we are going to refer to the membrane side where the NBDs are located as *in* (or *inside*), and the side of the membrane where the TMDs protrude into as *out* (or *outside*)^3^.

Hydrolysis of ATP and nucleotide exchange drive conformational changes that switch the arrangement of the substrate binding site on the two sides of the membrane. When bound to ATP, the TMDs are ready to catch substrates from, or release them to, the *out* side of the membrane. This conformation is usually called *open-outside*. Conversely, when bound to ADP the TMDs are *open-inside*, exchanging substrates with the *in* side of the membrane. The presence of a bound substrate has a strong impact on ATP hydrolysis or nucleotide exchange rates (or both) typically resulting in an acceleration of the ATPase cycle of ABC transporters^9–12^. Since the bound substrate does not directly contact the NBD, it must affect nucleotide processing (hydrolysis and/or exchange) through long-distance, allosteric conformational changes^13, 14^. Thus, the ATP-bound state must by necessity be an ensemble of at least two different conformations, characterized by different nucleotide-processing rates, and whose relative equilibrium is tuned by substrate binding so that it can modulate the observed ATPase rate.

We capture these basic features through the model depicted in Fig. 2A. The *open-outside*, ATP-bound state is split in two states: a slow-hydrolyzing/exchanging state T (TS when bound to substrate), and a fast-hydrolyzing/exchanging state T* (T*S when bound to substrate). The transition rates between T and T* (and also between TS and T*S), and hence their relative populations, can be modulated by the presence of a bound substrate. Indeed, the latter favors the fast nucleotide processing state, while the slow processing one is preferred when no substrate is bound. This mechanism allows for a control of the overall ATPase rate, as observed in experiments, through an indirect, allosteric switch operated by the substrate. For the sake of simplicity, the model proposed here considers that one single nucleotide affects the conformation of the transporter, although a more realistic description would capture the binding of two nucleotides possibly in their various ATP/ADP combinations.

**Figure 2.**
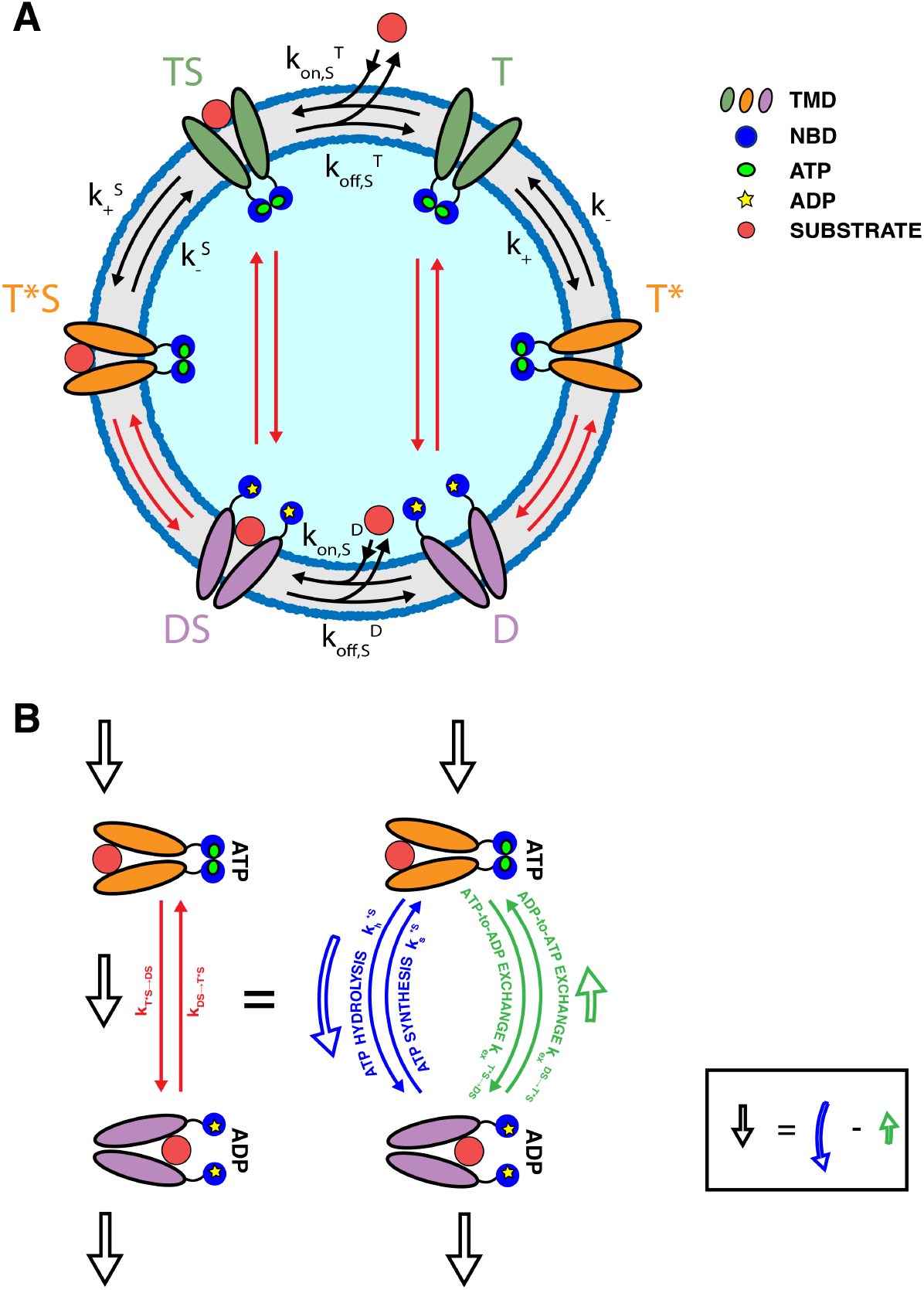
Model of ABC transporter. A) ABC transporters are embedded in a membrane, with the ATPase domains (blue circles) sitting on one side of it, and bind ATP (small green circles) or ADP (small yellow circles). Two different ATP-bound, open outside conformations (green or orange trans-membrane domains) are populated differently depending on the presence/absence of a bound substrate (orange circles). The ADP-bound, open-inside conformation (violet trans-membrane domains) is accessed from the ATP-bound conformations through composite reactions (red arrows) B) The composite reactions are simplified representations of reactions that can take place along different branches: ATP hydrolysis/synthesis (blue arrows) and ATP-to-ADP or ADP-to-ATP exchange; the overall flux through the reaction (black hollow arrow) is the difference between the fluxes taking place along the two branches, taken in opposite directions.

Upon hydrolysis or exchange, the ATP-bound *open-outside* states can convert into the ADP-bound *open-inside* conformation (D without, and DS with, bound substrate). Substrates can bind to the T and D, and unbind from the TS and DS, states. We simplify here the model by assuming that the T* state cannot directly bind substrates from solution, nor the T*S state can release them. Some special care must be paid to the transitions between states bound to different nucleotides, and to the associated rates (red arrows in Fig. 2A). These transitions can proceed via two different routes: exchange or hydrolysis/synthesis (Fig. 2B, green and blue arrows respectively). The rates of each individual route are related by relations imposed by thermodynamic equilibrium (see Supplementary Information Sec. A for a detailed derivation). In the absence of a bound substrate:

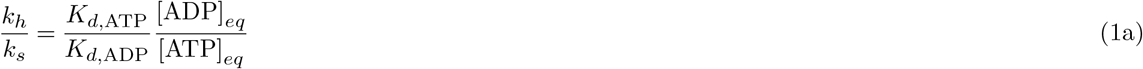

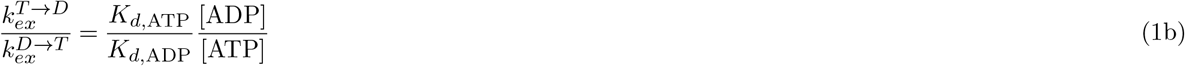

where *k*_*h*_ and *k*_*s*_ are the hydrolysis and synthesis rates, respectively, 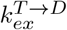 is the exchange rate for the release of ATP and binding of ADP, and 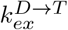 is the rate of the reverse exchange transition^15^. *K*_*d*,ATP_ and *K*_*d*,ADP_ are the dissociation constants of ATP and ADP respectively. Analogous relations hold in presence of the substrate. Each pair of transitions, namely T →D, T*→D, TS →DS and T*S →DS, supports its own rates and dissociation constants for the nucleotides, but always bound to each other by Eq. (1a) and (1b). These relations also imply that, away from equilibrium (*i*.*e*. [ATP]_*eq*_/[ADP]_*eq*_ ≠ [ATP]/[ADP]) the two branches of the reaction are not balanced and there is a net current that, flowing through the whole reaction network, can potentially result in a net current of molecules across the membrane. Hence, the substrate concentration difference across the membrane must be related to the energy available from ATP hydrolysis, Δ*G*:

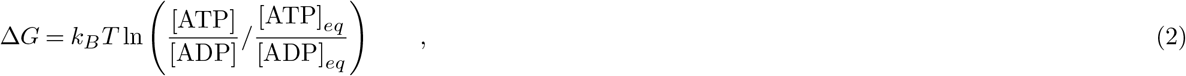

which vanishes at equilibrium. In Eq. (2) we have neglected the contribution from inorganic phosphate P_*i*_, assuming that its concentration is much larger than the one of nucleotides, as in the cell^16^, and is thus essentially unaltered by hydrolysis and synthesis.

All the rates of the system are additionally connected by further thermodynamic relations^17^, ensuring that if Δ*G* were zero, the system would satisfy detailed balance (see Supp. Info., Table S1). Hence, these thermodynamic constraints reduce the number of independent rates. As an important consequence, thermodynamic relations dictate that the binding of the substrate not only affects the ratio of the T and T* populations ([T*S]/[TS] = [T*]/[T]) as expected, but it must necessarily modify also their ATPase rates.

### The emerging logic of ABC transporters

In keeping with the role of ABC transporters in shuttling substrates against a concentration gradient across a membrane, we look at the steady-state ratio between the substrate concentration on the two sides of the membrane, [*in*]/[*out*], which is the most natural quantity to describe the functioning of the transporter.

The steady-state solution of the set of rate equations governing the system can be cast in a very instructive form (see Methods, and Supp. Info. Sections B and C):

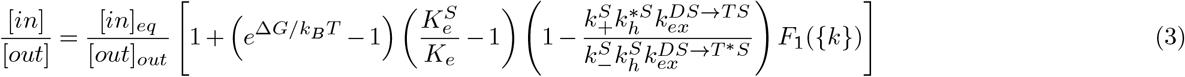

where *K*_*e*_ = *k*_+_/*k*_−_, and 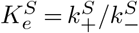, *i*.*e*. the equilibrium constants between T and T*, and TS and T*S, respectively. Here, *F*_1_({*k*}) is a positive function of the rates, whose specific expression is given in the Supplementary Information, Eq. (S19). The equilibrium concentration gradient is:

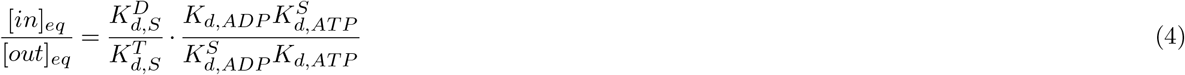

where 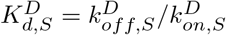 is the dissociation constant of the complex of the substrate with the ADP-bound, open-inside conformation, and 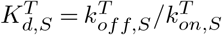 is the dissociation constant of the complex of the substrate with the ATP-bound, open-outside conformation of the transporter. *K*_*d*,ATP_ and *K*_*d*,ADP_ are the dissociation constants of ATP and ADP, respectively, for the transporter in the absence of a bound substrate (the same quantities for the substrate-bound transporter are indicated by a superscript *S*).

Eq. (3) reveals the intimate connection between ABC transporters and Maxwell Demons. In fact, it exhibits the structure of a logical “AND” function for the necessary requirements to let ABC transporters work (each one of them is highlighted by round brackets in the equation for the sake of clarity). Indeed, [*in*]/[*out*] reduces to the equilibrium value (Eq. (4)) when the following three conditions are not simultaneously satisfied:

i. Δ*G* ≠ 0: energy must be available from ATP hydrolysis. Without an energy source, it is impossible to push molecules against a concentration gradient, and the system falls back to equilibrium.
ii. 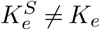: the relative populations of the two ATP-bound states must be tuned by substrate binding. In the absence of the substrate, the transporter preferentially sits in one state (T or T*), and switches to the other in the presence of a substrate (TS or T*S). In the language of the Maxwell Demon, the *measurement* corresponds to substrate binding, which is followed by the first *feedback* step, that is the setting of the memory bit by allostery. The necessity for the two equilibrium constants to be different is then clear: if they were equal, the memory would not change state upon binding, that is, the Demon could not record the measurement outcome, making the subsequent steps immaterial.
iii. 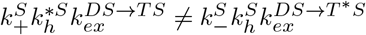: the ratio between these two terms quantifies the weight of the path to import a substrate with respect to the one of exporting a substrate. Transitions between states are always in principle reversible, and this is captured in the model introduced here. As a consequence, importers could in principle work as exporters and viceversa. The inequality between the two rate products ensures that one process dominates over the other by requiring that it proceeds in a definite order: the sequence of events comprising T →TS (substrate binding, or *measurement* in the Demon language), TS → T*S followed by T*S →DS (*feedback*: writing the outcome of the measurement and opening the door through hydrolysis), and DS →D followed by D →T, (*resetting* by substrate release and nucleotide exchange) cannot be equivalent to the opposite one. As a note, the precise import (export) path is immaterial for these considerations, provided the presence of one ATP hydrolysis step and one nucleotide exchange from ADP to ATP. In fact, using the relations enforced by thermodynamic consistency, the ratio between the product of the rates of all these paths is always captured by 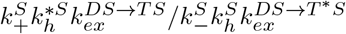 (see Supp. Info., Section D).

These three conditions exactly correspond to the three necessary requirements for the successful action of a Maxwell Demon: 1) it must consume energy; 2) the measurement (here, substrate binding) must be recorded on a physical storage device; 3) operating the door must depend on the gathered information, and the system must thus have a preferential direction for biochemical cycles. These three requirements must *all* be satisfied if the system has to be moved away from equilibrium, and the structure of Eq. (3) precisely captures the logical AND-like condition through a product of three factors, each of which must not vanish.

The solution of a model based only on basic structural and biochemical information has thus revealed that ABC transporters indeed *are* Maxwell Demons (technically, *autonomous* Maxwell Demons, where the demon and the door are blended into a single physical system^18^). The structure of Eq. (3) also reveals that reversing the sign of the triple product turns an importer ([*in*]>[*out*]) into an exporter ([*in*]<[*out*]), leaving many routes for nature to evolve one from the other, by tinkering with the transition rates.

### Details of the model tune the performance, while the logic is robust

The functional form of Eq. (3) states that the logic of the system does not depend on the choice of the parameters (rates) of the system, whose values can only decide how well transport takes place. Thus, to explore their role we numerically solved the model. For the sake of simplicity, we fixed some of the parameters: we chose 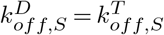 and 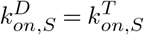, so that the substrate dissociation constants did not depend on the nucleotide state of the transporter, 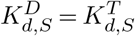. Moreover, we also set the nucleotide dissociation constants (both for ATP and ADP) to be independent on the presence or absence of a bound substrate, *i*.*e*. 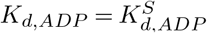 and 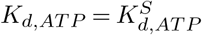. With these choices, from Eq. (4), [*in*]_*eq*_/[*out*]_*eq*_ = 1. Furthermore, we chose 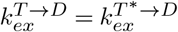 and 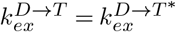, and analogously for the exchange rates for the substrate-bound transporter. With this simplification, and using the thermodynamic constraints on biochemical rates, condition (iii) from the previous section reduced to 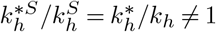. Finally we fixed the equilibrium constant between ATP and ADP, [*ATP*]_*eq*_/[*ADP*]_*eq*_ = 10^−9^, whereas in physiological condition [*ATP*]/[*ADP*] 10 ^16^ (see Methods, Table 1, for the values of rates used in simulations).

**Table 1.** Numerical values for the rates. The rates that are not defined in this table ensue from detailed balance conditions and thermodynamic constraints (see S.I., Table S1).

| Rate | Value | Rate | Value |
| --- | --- | --- | --- |
| $k_{\text{on},S}^T$ | $0.5 \mu\text{M}^{-1} \text{s}^{-1}$ | $k_{\text{off},S}^T$ | $0.01 \text{s}^{-1}$ |
| $k_{\text{on},S}^D$ | $0.5 \mu\text{M}^{-1} \text{s}^{-1}$ | $k_{\text{off},S}^D$ | $0.01 \text{s}^{-1}$ |
| $k_+$ | $1 \text{s}^{-1}$ | $k_+^S$ | $1 \text{s}^{-1}$ |
| $k_h$ | $0.02 \text{s}^{-1}$ | $k_h^S$ | $0.02 \text{s}^{-1}$ |
| | | $k_s^S$ | $2 \cdot 10^{-9} \text{s}^{-1}$ |
| $k_{-T}$ | $10^{-4} \text{s}^{-1}$ | $k_{-T}^S$ | $10^{-4} \text{s}^{-1}$ |
| $k_{-T}^*$ | $10^{-4} \text{s}^{-1}$ | $k_{-T}^{*S}$ | $10^{-4} \text{s}^{-1}$ |
| $k_{+T}$ | $0.5 \text{s}^{-1}$ | $k_{+T}^S$ | $0.5 \text{s}^{-1}$ |
| $k_{+T}^*$ | $0.5 \text{s}^{-1}$ | $k_{+T}^{*S}$ | $0.5 \text{s}^{-1}$ |
| $k_{+D}$ | $0.1 \text{s}^{-1}$ | $k_{+D}^S$ | $0.1 \text{s}^{-1}$ |
| $k_{+D}^*$ | $0.1 \text{s}^{-1}$ | $k_{+D}^{*S}$ | $0.1 \text{s}^{-1}$ |

We thus inspected, one by one, each of the necessary requirements:

i. Access to an energy source, Δ*G* ≠ 0. In equilibrium conditions, 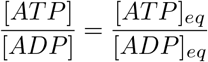, as predicted by Eq. (3), no transport is possible (Fig. 3A). For large values of the available energy, a plateau emerges, whose presence is due to rate-limiting steps that hinder the ability to fully convert the difference between the excess chemical potentials of ATP and ADP into a chemical potential difference across the membrane. The value of the plateau depends on all the parameters of the system.
ii. Physical underpinning of feedback 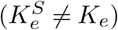. The plateau in Fig. 3A vanishes if 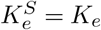, as predicted from the analytical solution, and becomes more pronounced as they become more different. This result signals that the transporter can best use the available energy when it can clearly tell the difference between the presence or absence of a substrate or, in a Maxwell Demon language, when the result of the measurement, to be used by the subsequent steps, can be stored more and more unambiguously. It is also noteworthy that as the ratio between the equilibrium constants changes across the unitary value, importers turning into exporters, as predicted in Eq. (3).
iii. Directionality of biochemical cycles 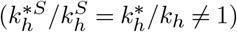. In typical cellular conditions (*α* = 10), Fig. 3B shows that 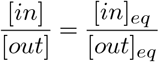 whenever 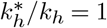, irrespective of all other rates, as expected. Furthermore, changing the ratio from being larger to smaller than 1, keeping all other rates fixed, changes importers into exporters, and vice versa. Also as expected, changing at the same time the inequalities for 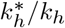 and 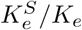 does not change the transport direction. The unitary value of the hydrolysis rates ratio corresponds thus to the perfect balance between inward and outward active, *i*.*e*. energy consuming, transport. The peculiar behavior for very large values of the 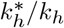 ratio, with the [*in*]/[*out*] ratio asymptotically moving back to equilibrium, is a consequence of the very large value of 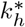 (since *k*_*h*_ is fixed for these figures) and of the corresponding synthesis rate (because of Eq. (1a)). Indeed, as 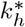 and 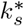 increase, the flux is more and more carried by the cycle T→TS→T*S→DS→D→T*→T always using the hydrolysis/synthesis reactions, which satisfies equilibrium thermodynamic constraints (see S.I., Eq. (S21))

**Figure 3.**
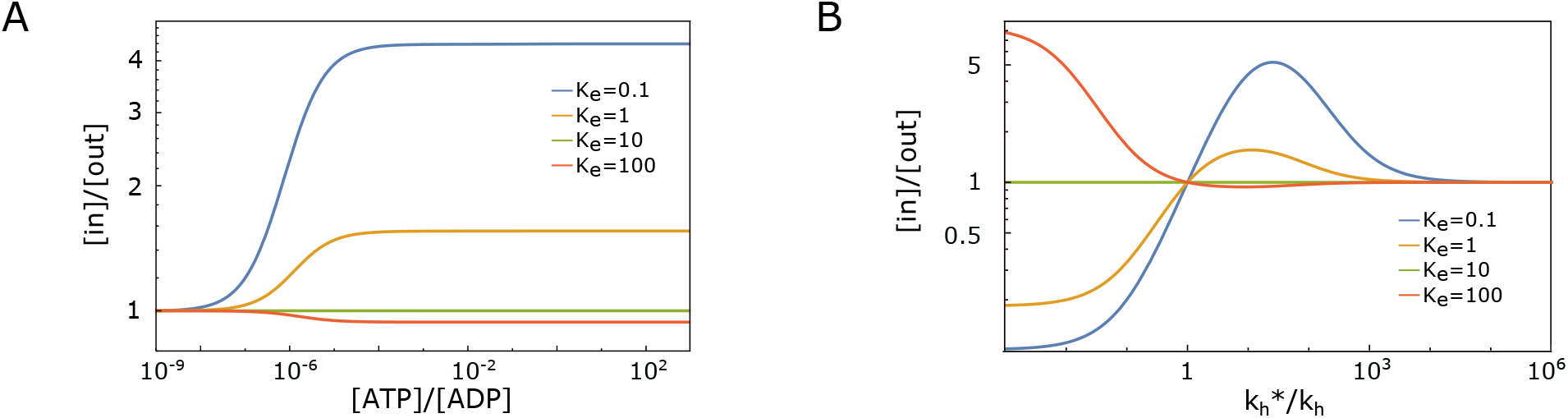
A) Steady state concentration ratio [*in*]/[*out*] as a function of the ratio [*ATP*]/[*ADP*], as a proxy for the available energy in the system. Results are shown for different values of *K*_*e*_, with 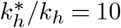 and 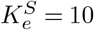. B) [*in*]/[*out*] as a function of the feedback efficiency on hydrolysis rates 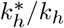. It is shown for 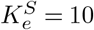 and different values of *K*_*e*_, in a far-from-equilibrium regime, with [*ATP*]/[*ADP*] = 10 and [*ATP*]_*eq*_/[*ADP*]_*eq*_ = 10^−9^, resembling typical cellular conditions.

The emerging *logic* of the model is actually robust upon inclusion of further details. For example, we could allow the T* conformation to bind/release substrates, with dissociation constant 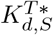. Thermodynamic consistency relations would then require 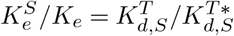, giving a clearer interpretation to the molecular stabilization of the T* conformation upon substrate binding. Considering thus also these new binding/unbinding processes led to a dependence of [*in*]/[*out*] on the absolute substrate concentrations, which was absent from the simpler model. In particular, substrate excess had here an inhibiting effect (Fig. 4A), as also observed in experiments ^19^. Nonetheless, [*in*]/[*out*] was shifted from its equilibrium value only if each of the three above conditions was respected, the dependence on the absolute concentrations thus restricted to the functional form of *F*_1_ in Eq. (3). These findings were to be expected, because, whereas the overall performance of the transporter might depend on the details of the model and on its associated parameters, its internal logic represents a hard set of rules.

**Figure 4.**
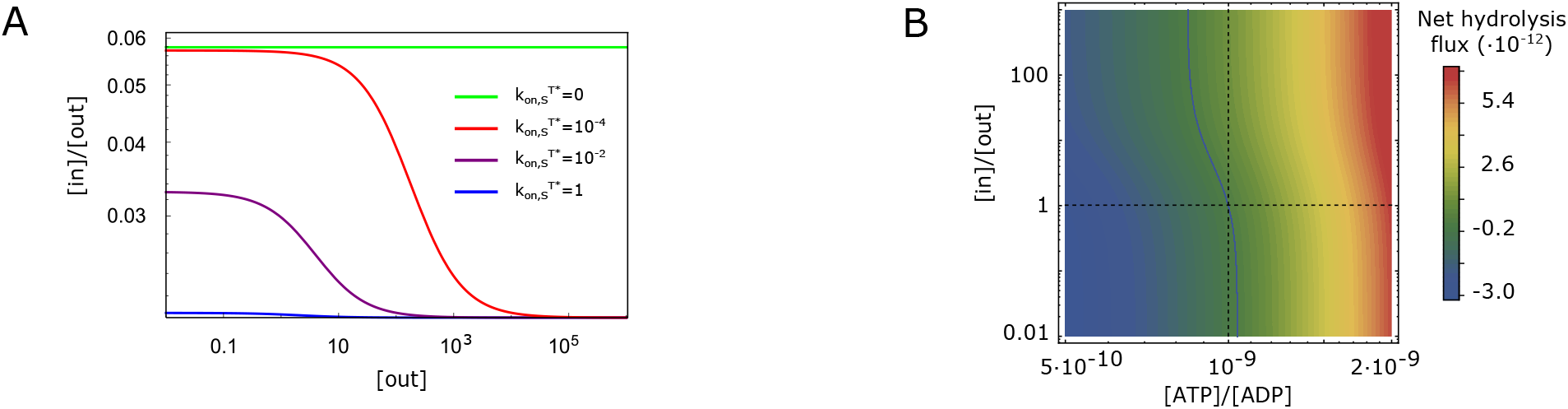
A) Steady-state ratio [*in*]/[*out*] as a function of substrate concentration outside the cell for different binding rates 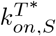 between T* and T*S. B) Net hydrolysis rate Φ_*h*_ in a close-to-equilibrium regime, when a substrate concentration gradient is imposed on the system. The horizontal and vertical dashed lines represent equilibrium conditions (respectively [*in*]/[*out*] = 1 and *α* = *α*_*eq*_ = 10^−9^). The continuous line corresponds to the limit when hydrolysis and synthesis fluxes are equal (Φ_*h*_ = 0).

The full kinetic reversibility of the present model allowed us to establish a connection between the available energy and transport. Indeed, had synthesis been neglected, the free-energy liberated by ATP hydrolysis would have diverged, positioning the system always in the plateau region of Fig. 3A. Considering synthesis explicitly, instead, made it possible to find conditions where there was a net production of ATP from ADP, typically in connection with a substrate flux across the transporter opposite to the one that would have been supported in physiological conditions, thus forcing importers to export molecules and vice versa. For example, this would happen in the presence of an excess of ADP over ATP beyond what their thermodynamic equilibrium would dictate and/or in the presence of a substrate ratio [*in*]/[*out*] in excess over the one that the system would have at steady-state, given the other conditions (Fig. 4B). This result is consistent with the detection of ATP synthesis from ADP by ABC exporters when forced to import substrates in the presence of excess ADP^20^.

### The cost of processing information

Besides its paradigmatic simplicity, the Maxwell Demon metaphor bridges between the microscopic, molecular description and the more abstract, but not less fundamental approach of information theory. Indeed, as previously mentioned, the solution to the thermodynamic paradox of the Maxwell Demon operation came precisely by considering the free-energy cost of information processing. The exact correspondence between ABC transporters and Maxwell Demons, thus, calls for a more direct approach in terms of information theory.

In particular, we expected that the excess chemical potential across the membrane

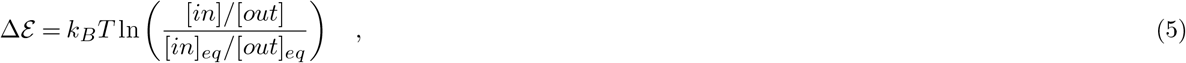

had to correspond to the available free-energy from ATP hydrolysis, Δ*G*, decreased by the free-energy associated to acquiring information (*measure*), using it (*feedback*) and erasing it (*reset*)

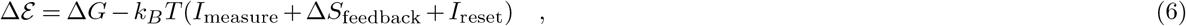

In the case of an importer, the measurement operation corresponds to substrate binding and to the consequent transition from the T to the TS state. The associated information gain is^21^

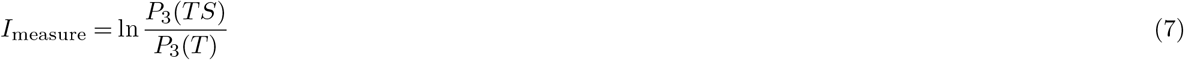

where *P*_3_(*T*) and *P*_3_(*TS*) are the stationary probabilities of the T state *conditional* to the absence or presence of a bound substrate, respectively. They must thus be computed on the corresponding restricted 3-state subsystems (T-T*-D and TS-T*S-DS) depicted in Fig. 5. A similar argument can be used for *I*_reset_ which corresponds to the release of the substrate, *I*_reset_ = ln *P*_3_(*D*)*/P*_3_(*DS*).

**Figure 5.**
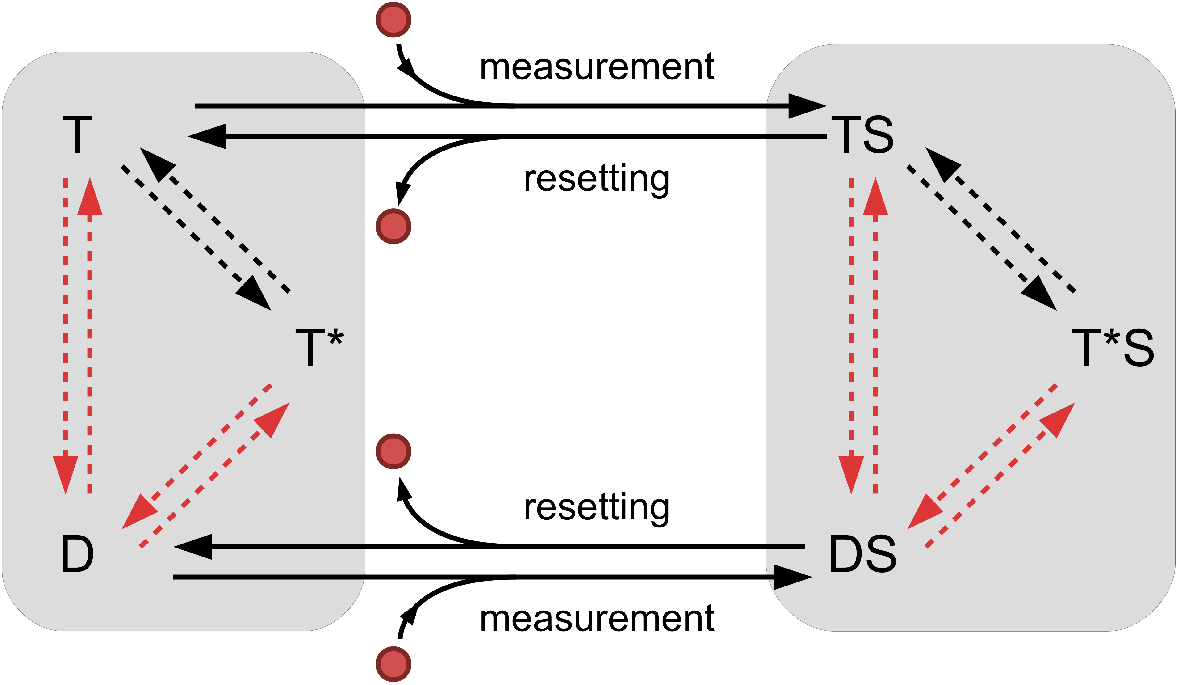
Each of the grey rectangles corresponds to a three-state system either with (TS-T*S-DS) or without (T-T*-D) bound substrate. Solid arrows correspond to measurement and resetting processes (respectively materialized by binding and unbinding of substrate). Dashed arrows are the rates realizing the feedback. As in Fig.2, red arrows identify composite reactions.

Δ*S*_feedback_ is the entropy dissipated into the environment (thus heat from the relation Δ*S* = Δ*Q/T*) by the actuation of the feedback mechanism. An ABC importer transports substrates across the membrane through ATPase cycles in which both hydrolysis and nucleotide exchange are used (all other cycles always respect equilibrium, according to thermodynamic constraints). In particular, energy is exploited hydrolyzing ATP when the substrate is bound (after measure), and exchanging nucleotide when the substrate is unbound (after reset), so that the system is ready to measure again and restart the cycle. As expected, independently of the specific cycle considered, as long as it runs according to this rule, it dissipates (see Supp. Info., Section E)^22^

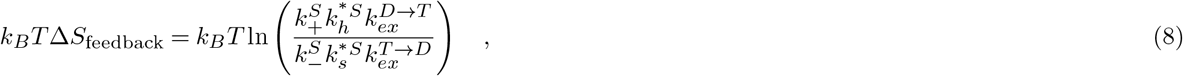

which is precisely the net heat released into the environment by opening the door after measure and closing it after the molecule has passed 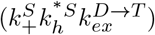 rather than going through the opposite process 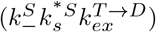.

With these ingredients, it is possible to show that Eq. (6) coincides with Eq. (3) (by taking the logarithm of both sides), showing that starting from an information-theoretic approach immediately provides the correct structure of the solution, to be then made explicit using the biochemical rates.

## Discussion

Transport against a concentration gradient is a crucial process for cells, both to import nutrients, to export metabolic products or other noxious molecules (toxins, antibiotics etc.) and to shuttle molecules across internal cellular membranes. ABC transporters, one of the major classes of proteins that carry out this task, have been fairly well characterized both structurally and biochemically. The picture that has emerged bears tantalizing similarities with the Maxwell Demon, which is an idealised device able to shift the steady-state of a system from its equilibrium prescription. In this work we have built on these sources of information to propose a simple model of ABC transporters whose solution shows that they are not simply analogous to Maxwell Demons: they *are* Maxwell Demons. Already billions of years ago, evolution has played with the different conformations and their transition rates to write in the system the precise necessary conditions for the realization of a Maxwell Demon.

Without any doubt our model, which is on purpose as simple as possible, omits several details of the structure and biochemistry of ABC transporters: two ATP molecules are likely to be hydrolysed for each transported molecule; the products of ATP hydrolysis are ADP and inorganic phosphate (Pi), which are then sequentially released leading to further sub-conformations; nucleotide exchange might proceed through a fully apo-state (no nucleotides) or through mixed ATP/ADP bound states. Nonetheless, our work strongly suggests that, even within a more complex but more realistic scheme, it would still be necessary to identify the basic steps of a Maxwell Demon: measurement, feedback and resetting. Indeed, the logical structure pinpointing the necessary ingredients for the functioning of ABC transporters can be directly derived from an information-theoretic approach.

From a biological perspective, besides the basic essential requirements for the transporter to work, much freedom remains in the choice of the parameter values for its optimisation, according to a plethora of criteria, which might not be simply the maximisation of the gradient across the membrane. Furthermore, our model shows that there are several different routes to turn an importer into an exporter, leaving much freedom to evolve one into the other.

Our conclusions here have further broader consequences for any molecular machine able to bring the system it acts upon into a non-equilibrium steady state. Not only it needs an external energy source. It also needs obligatorily a “measurement device”, that is, a way to distinguish between different substrates, likely through a direct interaction that, allosterically, modulates the populations of at least two different conformations, thus writing the measurement result in a sort of *molecular information bit*. Furthermore, it is mandatory that upon interaction, a specific feedback action takes place, which is possible if, for example, the different conformations selected by the substrate allow the energy-consuming cycle to proceed at different rates. These are features that any structural/biochemical investigation must address, together with careful measurements of the kinetic rates of the system, rather than only of its thermodynamic properties (*i*.*e*. equilibrium constants and dissociation rates).

From a broader perspective, our work also suggests that the language of information theory, which is often advocated as the most suitable framework to deal with biological systems beyond the molecular details^23^, can be derived as an *emerging feature* from a reductionist approach based on biophysical and biochemical insight.

## Methods

### Derivation of the stationary concentration gradient

The system evolves according to a Master Equation. The concentrations of the six possible states (*T, T*^*^, *D, TS, T*^*^*S*, and *DS*) satisfy six coupled rate equations, detailed in the Supplementary Information, Sections A and B. In the Supp. Info., Section A, we also introduce the correct expressions for composite reactions. We must add to these six equations the normalization condition, i.e. the conservation of the total concentrations of transporters: [*T*] +[*T*^*^] +[*D*] +[*TS*] +[*T*^*^*S*] +[*DS*] = *C*_tot_. The probability to be in a given state, *X*, is *P*(*X*) = [*X*]*/C*_tot_, and its value at stationarity is *P*^st^(*X*). The stationary solution is derived explicitly in the Supp. Info., Section B.

Moreover, at stationarity, there must be a steady rate of binding and unbinding of substrates, which translates into the absence of a net flux both between *T* and *TS*, and between *D* and *DS*. This condition is:

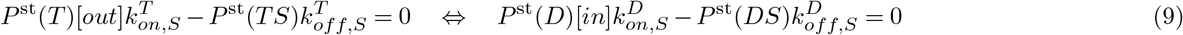

We introduce the following notation:

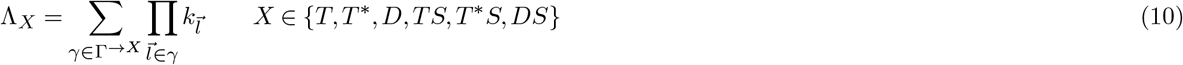

where Γ is the set of all spanning trees directed towards *X, γ* an element of this set, and 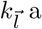 a rate belonging to *γ*, with the correct orientation. We finally have the following expression for the stationary concentration gradient^24^:

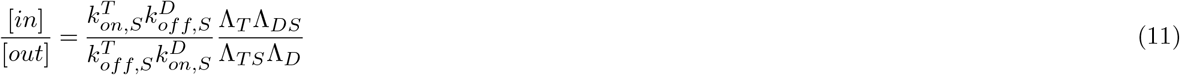

This is manifestly equal to Eq. (6), since any ratio Λ(*X*)/Λ(*Y*) = *P*_3_(*X*)*/P*_3_(*Y*), where *P*_3_(*X*) is the stationary probability to be in the state *X*, evaluated as only the three-state subsystem which *X* belongs to, namely (*T*−*T*^*^− *D*) or (*TS*−*T*^*^*S*−*DS*), does exist. Eq. (11) can be further manipulated to obtain Eq. (3) of the main text, as discussed in the Supp. Info., Section C.

### Numerical values

If not stated otherwise, the numerical values used for the simulations are shown in table 1.

## Acknowledgements

S. Flatt thanks the Swiss National Science Foundation for support through grant 200020_178763.

## Author contributions statement

All authors conceived the model. S.F. performed analytical and numerical calculations. All authors wrote and reviewed the manuscript.

## Additional information

**Competing interests** The authors declare no competing interests.

