## Supplemental material for "ABC Transporters are billion-year-old Maxwell Demons"

### A The model

As illustrated in Figure 2B, the rates from the state T to the state D, as well as the ones connecting T* and D, TS and DS, and T*S and DS are composite reactions constituted by hydrolysis/synthesis and exchange processes. The rates are:

$$k_{T\to D} = k_h + k_{ex}^{T\to D} \qquad k_{D\to T} = k_s + k_{ex}^{D\to T} \tag{S1}$$

$$k_{T^*\to D} = k_h^* + k_{ex}^{T^*\to D} \qquad k_{D\to T^*} = k_s^* + k_{ex}^{D\to T^*} \tag{S2}$$

$$k_{TS\to DS} = k_h^S + k_{ex}^{TS\to DS} \qquad k_{DS\to TS} = k_s^S + k_{ex}^{DS\to TS} \tag{S3}$$

$$k_{T^*S\to DS} = k_h^{*S} + k_{ex}^{T^*S\to DS} \qquad k_{DS\to T^*S} = k_s^{*S} + k_{ex}^{DS\to T^*S} \tag{S4}$$

Exchange processes correspond to the unbinding of one nucleotide(ATP or ADP), with the subsequent binding of the other (ADP or ATP, respectively). Hence, the rates are[1]

$$\begin{aligned}
k_{ex}^{D\to T} &= k_{-ADP}\frac{[ATP]k_{+ATP}}{[ATP]k_{+ATP}+[ADP]k_{+ADP}} & k_{ex}^{T\to D} &= k_{-ATP}\frac{[ADP]k_{+ADP}}{[ATP]k_{+ATP}+[ADP]k_{+ADP}}\\
k_{ex}^{D\to T^*} &= k^*_{-ADP}\frac{[ATP]k^*_{+ATP}}{[ATP]k^*_{+ATP}+[ADP]k^*_{+ADP}} & k_{ex}^{T^*\to D} &= k^*_{-ATP}\frac{[ADP]k^*_{+ADP}}{[ATP]k^*_{+ATP}+[ADP]k^*_{+ADP}}\\
k_{ex}^{DS\to TS} &= k^S_{-ADP}\frac{[ATP]k^S_{+ATP}}{[ATP]k^S_{+ATP}+[ADP]k^S_{+ADP}} & k_{ex}^{TS\to DS} &= k^{*S}_{-ATP}\frac{[ADP]k^{*S}_{+ADP}}{[ATP]k^{*S}_{+ATP}+[ADP]k^{*S}_{+ADP}}\\
k_{ex}^{DS\to T^*S} &= k^{*S}_{-ADP}\frac{[ATP]k^{*S}_{+ATP}}{[ATP]k^{*S}_{+ATP}+[ADP]k^{*S}_{+ADP}} & k_{ex}^{T^*S\to DS} &= k^{*S}_{-ATP}\frac{[ADP]k^{*S}_{+ADP}}{[ATP]k^{*S}_{+ATP}+[ADP]k^{*S}_{+ADP}}
\end{aligned} \tag{S5}$$

It follows that the ratio of two reverse exchange rates is given by Eq. 1b (and the same for the different superscripts associated to the states $TS$, $T^*$ and $T^*S$).

All the rates in Figures 2A and 2B must satisfy thermodynamic constrains at equilibrium. In particular, all cycles have to fulfill the Kolmogorov condition, which corresponds to the requirement of detailed balance. Thus, not all rates are independent: as an example the constraint between hydrolysis-synthesis and exchange is given by

$$\left.\frac{k_h k_{ex}^{D\to T}}{k_s k_{ex}^{T\to D}}\right|_{eq} = 1 \quad , \tag{S6}$$

hence leading to Eq. 1a.

In Table S1 we list all the rates as a function of independent system parameters.

For the sake of clarity and until the end of this section, we introduce the parameters $\alpha := [ATP]/[ADP]$ and $\eta := k_h^*/k_h = k_h^{*S}/k_h^S$.

**Table S1.** Expression of the rates as a function of independant parameters

| **Rate** | **Expression** |
|---|---|
| $k^T_{on,S}$ | $k^T_{on,S}$ |
| $k^D_{on,S}$ | $k^D_{on,S}$ |
| $k_+$ | $k_+$ |
| $k_-$ | $\frac{k_+}{K_e}$ |
| $k_h$ | $k_h$ |
| $k_{ex}^{T\to D}$ | $\frac{k_{-T}k_{+D}}{k_{+D}+\alpha k_{+T}}$ |
| $k_h^*$ | $\eta k_h$ |
| $k_{ex}^{T^*\to D}$ | $\frac{k^*_{-T}k^*_{+D}}{k^*_{+D}+\alpha k^*_{+T}}$ |
| $k_h^S$ | $k_h^S$ |
| $k_{ex}^{TS\to DS}$ | $\frac{k^S_{-T}k^S_{+D}}{k^S_{+D}+\alpha k^S_{+T}}$ |
| $k_h^{*S}$ | $\eta k_h^S$ |
| $k_{ex}^{T^*S\to DS}$ | $\frac{k^{*S}_{-T}k^{*S}_{+D}}{k^{*S}_{+D}+\alpha k^{*S}_{+T}}$ |

| **Rate** | **Expression** |
|---|---|
| $k^T_{off,S}$ | $k^T_{off,S}$ |
| $k^D_{off,S}$ | $k^D_{off,S}$ |
| $k_+^S$ | $k_+^S$ |
| $k_-^S$ | $\frac{k_+^S}{K_e^S}$ |
| $k_s$ | $\frac{[in]_{eq}}{[out]_{eq}}\frac{k_h k^T_{off,S}k^D_{on,S}k^S_s}{k^S_h k^D_{off,S}k^T_{on,S}}$ |
| $k_{ex}^{D\to T}$ | $\frac{[in]_{eq}}{[out]_{eq}}\frac{\alpha k_{-T}k^T_{off,S}k^D_{on,S}k_{+D}k^S_s}{\alpha_{eq}k^S_h k^D_{off,S}k^T_{on,S}(k_{+D}+\alpha k_{+T})}$ |
| $k_s^*$ | $\frac{[in]_{eq}}{[out]_{eq}}\frac{\eta K_e k_h k^T_{off,S}k^D_{on,S}k^S_s}{k^S_h k^D_{off,S}k^T_{on,S}}$ |
| $k_{ex}^{D\to T^*}$ | $\frac{[in]_{eq}}{[out]_{eq}}\frac{\alpha K_e k^*_{-T}k^T_{off,S}k^D_{on,S}k^*_{+D}k^S_s}{\alpha_{eq}k^S_h k^D_{off,S}k^T_{on,S}(k^*_{+D}+\alpha k^*_{+T})}$ |
| $k_s^S$ | $k_s^S$ |
| $k_{ex}^{DS\to TS}$ | $\frac{\alpha k^S_{-T}k^S_{+D}k^S_s}{\alpha_{eq}k^S_h(k^S_{+D}+\alpha k^S_{+T})}$ |
| $k_s^{*S}$ | $\eta K_e^S k_s^S$ |
| $k_{ex}^{DS\to T^*S}$ | $\frac{\alpha K^S_e k^{*S}_{-T}k^{*S}_{+D}k^S_s}{\alpha_{eq}k^S_h(k^{*S}_{+D}+\alpha k^{*S}_{+T})}$ |

Expression of the rates as a function of independent parameters, after introducing detailed balance constrains as well as $\eta, K_e$ and $K_e^S$. All the terms in the right column "Expression" can be defined independently.

### B Derivation of the steady-state of the system

The evolution of the system is described by the following Master Equation:

$$\begin{cases} \dfrac{\mathrm{d}[TS]}{\mathrm{d}t} = -[TS](k_+^S + k_{TS\to DS} + k_{off,S}^T) + [T^*S]k_-^S + [DS]k_{DS\to TS} + [T][out]k_{on,S}^T & \text{(S7a)} \\ \dfrac{\mathrm{d}[T^*S]}{\mathrm{d}t} = [TS]k_+^S - [T^*S](k_-^S + k_{T^*S\to DS}) + [DS]k_{DS\to T^*S} & \text{(S7b)} \\ \dfrac{\mathrm{d}[DS]}{\mathrm{d}t} = [TS]k_{TS\to DS} + [T^*S]k_{T^*S\to DS} - [DS](k_{DS\to TS} + k_{DS\to T^*S} + k_{off,S}^D) + [D][in]k_{on,S}^D & \text{(S7c)} \\ \dfrac{\mathrm{d}[T]}{\mathrm{d}t} = [TS]k_{off,S}^T - [T]([out]k_{on,S}^T + k_+ + k_{T\to D}) + [T^*]k_- + [D]k_{D\to T} & \text{(S7d)} \\ \dfrac{\mathrm{d}[T^*]}{\mathrm{d}t} = [T]k_- [T^*](k_- + k_{T^*\to D}) + [D]k_{D\to T^*} & \text{(S7e)} \\ \dfrac{\mathrm{d}[D]}{\mathrm{d}t} = [DS]k_{off,S}^D + [T]k_{T\to D} + [T^*]k_{T^*\to D} - [D]([in]k_{on,S}^D + k_{D\to T} + k_{D\to T^*}) & \text{(S7f)} \end{cases}$$

Note that we keep the short notation for composite reactions defined in the S.I. Section A. To the set of equations above, we must add the normalization condition, that is the conservation of the total concentration of transporters:

$$[TS] + [T^*S] + [DS] + [T] + [T^*] + [D] = C_{\mathrm{tot}} \tag{S8}$$

so that the probability to be in the state $X$ is $P(X) = [X]/C_{\mathrm{tot}}$. At steady state, all time derivatives are equal to zero. Naming the stationary probability to be in $X$ as $P^{\mathrm{st}}(X)$, and introducing the following notation

$$\Lambda_X = \sum_{\gamma\in\Gamma^{\to X}} \prod_{\vec{l}\in\gamma} k_{\vec{l}} \qquad X \in \{T, T^*, D, TS, T^*S, DS\} \tag{S9}$$

$$\Psi_{T^*} = k_- + k_{T^*\to D} \quad \text{and} \quad \Psi_{T^*S} = k_-^S + k_{T^*S\to DS} \tag{S10}$$

as described in the Methods, the solution can be written in a compact way (with a proportionality factor accounting for the normalization of the probability distribution)[2]

$$\begin{aligned} P^{\mathrm{st}}(TS) &\propto \Lambda_{TS}[out]k_{on,S}^T\Lambda_T + \Lambda_{TS}[out]k_{on,S}^T[in]k_{on,S}^D\Psi_{T^*} + \Psi_{T^*S}[out]k_{on,S}^T k_{off,S}^D\Lambda_T + \Lambda_{TS}[in]k_{on,S}^D\Lambda_D \\ P^{\mathrm{st}}(T^*S) &\propto \Lambda_{T^*S}[out]k_{on,S}^T\Lambda_T + \Lambda_{T^*S}[in]k_{on,S}^D\Lambda_D + \Lambda_{T^*S}[out]k_{on,S}^T[in]k_{on,S}^D\Psi_{T^*} + k_+^S[out]k_{on,S}^T k_{off,S}^D\Lambda_T + k_{DS\to T^*S}[in]k_{on,S}^D k_{off,S}^T\Lambda_D \\ P^{\mathrm{st}}(DS) &\propto \Lambda_{DS}\Lambda_D[in]k_{on,S}^D + \Lambda_{DS}[outk_{on,S}^T[in]k_{on,S}^D\Psi_{T^*} + \Lambda_D k_{off,S}^T + [in]k_{on,S}^D\Psi_{T^*S} + \Lambda_T[out]k_{on,S}^T\Lambda_{DS} \\ P^{\mathrm{st}}(T) &\propto \Lambda_{TS}k_{off,S}^T\Lambda_T + \Lambda_T k_{off,S}^T k_{off,S}^D\Psi_{T^*S} + \Lambda_{TS}[in]k_{on,S}^D k_{off,S}^T\Psi_{T^*} + \Lambda_{DS}k_{off,S}^D\Lambda_T \\ P^{\mathrm{st}}(T^*) &\propto \Lambda_{T^*}k_{off,S}^T\Lambda_{TS} + \Lambda_{T^*}k_{off,S}^D\Lambda_{DS} + \Lambda_{T^*}k_{off,S}^T k_{off,S}^D\Psi_{T^*S} + k_+ k_{off,S}^T[in]k_{on,S}^D\Lambda_{TS} + k_{D\to T^*}k_{off,S}^D[out]k_{on,S}^T\Lambda_{DS} \\ P^{\mathrm{st}}(D) &\propto \Lambda_{DS}k_{off,S}^D\Lambda_D + \Lambda_D k_{off,S}^D k_{off,S}^T\Psi_{T^*S} + \Lambda_{DS}[out]k_{on,S}^T k_{off,S}^D\Psi_{T^*} + \Lambda_{TS}k_{off,S}^T\Lambda_D \end{aligned} \tag{S11}$$

Finally, we employ the condition that at stationarity there must be a steady rate of binding/unbinding, that is the absence of net flux of substrates both between T and TS, and D and DS. This corresponds to:

$$P^{\mathrm{st}}(T)[out]k_{on,S}^T - P^{\mathrm{st}}(TS)k_{off,S}^T = 0 \quad \Leftrightarrow \quad P^{\mathrm{st}}(D)[in]k_{on,S}^D - P^{\mathrm{st}}(DS)k_{off,S}^D = 0 \tag{S12}$$

The substitution of the expressions for $P^{\mathrm{st}}(TS)$ and $P^{\mathrm{st}}(T)$ (S11) into (S12) leads to

$$\begin{aligned} &\left(\Lambda_{TS}k_{off,S}^T\Lambda_T + \Lambda_T k_{off,S}^T k_{off,S}^D\Psi_{T^*S} + \Lambda_{TS}[in]k_{on,S}^D k_{off,S}^T\Psi_{T^*} + \Lambda_{DS}k_{off,S}^D\Lambda_T\right)\cdot[out]k_{on,S}^T \\ -\ &\left(\Lambda_{TS}[out]k_{on,S}^T\Lambda_T + \Lambda_{TS}[out]k_{on,S}^T[in]k_{on,S}^D\Psi_{T^*} + \Psi_{T^*S}[out]k_{on,S}^T k_{off,S}^D\Lambda_T + \Lambda_{TS}[in]k_{on,S}^D\Lambda_D\right)k_{off,S}^T = 0 \end{aligned} \tag{S13}$$

and finally:

$$\frac{[in]}{[out]} = \frac{k_{on,S}^T k_{off,S}^D}{k_{off,S}^T k_{on,S}^D}\frac{\Lambda_T\Lambda_{DS}}{\Lambda_{TS}\Lambda_D} \tag{S14}$$

### C Derivation and explicit expression of Eq. (3)

We will sketch the analytical development to obtain Eq. (3) from Eq. (S14). Introducing the symmetry constraints as well as the assymmetry induced by $K_e$ and $K_e^S$, we can rewrite Eq. (S14) as:

$$\frac{[in]}{[out]} = \frac{[in]_{eq}}{[out]_{eq}} \cdot \frac{\left[\Lambda_D + \left(\frac{K_e}{K_e^S} - 1\right) k_- k_{T\to D}\right] \Lambda_T}{\left[\Lambda_T + \left(\frac{K_e}{K_e^S} - 1\right) k_- k_{D\to T}\right] \Lambda_D} \tag{S15}$$

Thus, the displacement from equilibrium is given by

$$\frac{[in]/[out]}{[in]_{eq}/[out]_{eq}} - 1 = \frac{\frac{K_e}{K_e^S} - 1}{\left[\Lambda_T + \left(\frac{K_e}{K_e^S} - 1\right) k_- k_{D\to T}\right] \Lambda_D} \cdot k_- \cdot \left(k_- k_{T\to D} k_{D\to T^*} - k_+ k_{T^*\to D} k_{DtoT}\right) \tag{S16}$$

Splitting the total rates between $T$ and $D$ states (and similarly for their analogs) as a sum of hydrolyis/synthesis and exchange leads to:

$$\frac{[in]/[out]}{[in]_{eq}/[out]_{eq}} - 1 = \frac{\left(\frac{K_e}{K_e^S} - 1\right)\left(\frac{\alpha}{\alpha_{eq}} - 1\right)}{\left[\Lambda_T + \left(\frac{K_e}{K_e^S} - 1\right) k_- k_{D\to T}\right] \Lambda_D} \cdot k_- \cdot \left(k_s k_+ k_{ex}^{T^*\to D} - \frac{\alpha_{eq}}{\alpha} k_{ex}^{D\to T} k_+ k_h^*\right) \tag{S17}$$

Finally

$$\frac{[in]/[out]}{[in]_{eq}/[out]_{eq}} - 1 = \frac{\left(\frac{K_e}{K_e^S} - 1\right)\left(\frac{\alpha}{\alpha_{eq}} - 1\right)\left(1 - \frac{k_h^* k_{ex}^{T\to D}}{k_h k_{ex}^{T^*\to D}}\right)}{\left[\Lambda_T + \left(\frac{K_e}{K_e^S} - 1\right) k_- k_{D\to T}\right] \Lambda_D} \cdot k_- k_+ k_s^S k_{ex}^{T^*\to D} \tag{S18}$$

After a few rearrangements, the final result can be written in a more convenient way, with the with the explicit form for $F_1(\{k\})$ (Eq. (3)):

$$\frac{[in]}{[out]} = \frac{[in]_{eq}}{[out]_{eq}} \left[1 + \left(e^{\Delta G/k_B T} - 1\right)\left(\frac{K_e^S}{K_e} - 1\right)\left(1 - \frac{k_+^S k_h^{*S} k_{DS\to TS}}{k_-^S k_h^S k_{DS\to T^*S}}\right) \cdot \underbrace{\left(\frac{k_+^2 \cdot k_s^S \cdot k_{ex}^{T^*\to D}}{K_e \cdot \Lambda_{TS} \Lambda_D}\right)}_{F_1(\{k\})}\right] \tag{S19}$$

### D Ratio between import and export cycles

In the discussion of Eq. (3), we stress the role played by the ratio $\frac{k_+^S k_h^{*S} k_{ex}^{DS\to TS}}{k_-^S k_h^S k_{ex}^{DS\to T^*S}}$. This corresponds to the ratio between import and export paths going through one hydrolysis and one exchange. There are two such possible paths that move the system out of equilibrium:

$$\begin{aligned}
&\frac{T \to TS \to T^*S \xrightarrow{\text{hydrolysis}} DS \to D \xrightarrow{\text{exchange}} T}{T \xrightarrow{\text{hydrolysis}} D \to DS \xrightarrow{\text{exchange}} T^*S \to TS \to T} : \frac{k_{on,S}^T k_+^S k_h^{*S} k_{off,S}^D k_{ex}^{D\to T}}{k_h k_{on,S}^D k_{ex}^{DS\to T^*S} k_-^S k_{off,S}^T} = \frac{[in]_{eq}}{[out]_{eq}} \frac{k_+^S k_h^{*S} k_{ex}^{DS\to TS}}{k_-^S k_h^S k_{ex}^{DS\to T^*S}} \\
&\frac{T \to TS \xrightarrow{\text{hydrolysis}} DS \to D \xrightarrow{\text{exchange}} T^* \to T}{T \to T^* \xrightarrow{\text{hydrolysis}} D \to DS \xrightarrow{\text{exchange}} TS \to T} : \frac{k_{on,S}^T k_h^S k_{off,S}^D k_{ex}^{D\to T^*} k_-}{k_+ k_h^* k_{on,S}^D k_{ex}^{DS\to TS} k_{off,S}^T} = \frac{[in]_{eq}}{[out]_{eq}} \frac{k_+^S k_h^{*S} k_{ex}^{DS\to TS}}{k_-^S k_h^S k_{ex}^{DS\to T^*S}}
\end{aligned} \tag{S20}$$

However, if on both sides the transport cycle goes through the states $T^*S$ and $T^*$ the symmetry of the system brings the system back to equilibrium. The same result holds for a cycle that avoids the states $T^*S$ and $T^*$, only going through the four *central* states.

$$\begin{aligned}
&\frac{T \to TS \to T^*S \xrightarrow{\text{hydrolysis}} DS \to D \xrightarrow{\text{exchange}} T^* \to T}{T \to T^* \xrightarrow{\text{hydrolysis}} D \to DS \xrightarrow{\text{exchange}} T^*S \to TS \to T} : \frac{k_{on,S}^T k_+^S k_h^{*S} k_{off,S}^D k_{ex}^{D\to T^*} k_-}{k_+ k_h^* k_{on,S}^D k_{ex}^{DS\to T^*S} k_-^S k_{off,S}^T} = \frac{[in]_{eq}}{[out]_{eq}} \\
&\frac{T \to TS \xrightarrow{\text{hydrolysis}} DS \to D \xrightarrow{\text{exchange}} T}{T \xrightarrow{\text{hydrolysis}} D \to DS \xrightarrow{\text{exchange}} TS \to T} : \frac{k_{on,S}^T k_h^S k_{off,S}^D k_{ex}^{D\to T}}{k_h k_{on,S}^D k_{ex}^{DS\to TS} k_{off,S}^T} = \frac{[in]_{eq}}{[out]_{eq}}
\end{aligned} \tag{S21}$$

### E Heat released by feedback operations

Here, we show that all cycles involving hydrolysis after measurement, and exchange processes after resetting exhibit the same dissipation. Indeed, there are four different possible sets of feedback operations, each one associated with a certain heat release into the environment, $\Delta S$ [3]:

a) $TS \rightarrow T^*S \xrightarrow{\text{hydrolysis}} DS$, after measurement, i.e. with bound substrate $S$; $D \xrightarrow{\text{exchange}} T$, after resetting, i.e. without bound substrate. To compute the heat release, we employ the thermodynamic relations presented in Table 1. Hence, we have:

$$\frac{\Delta S}{k_B T} = \ln\left(\frac{k_+^S k_h^{*S} k_{ex}^{D\to T}}{k_-^S k_s^{*S} k_{ex}^{T\to D}}\right) = \ln\left(K_e^S \frac{k_h^{*S}}{k_s^{*S}} \frac{\alpha}{\alpha_{eq}} \frac{[in]_{eq}}{[out]_{eq}} \frac{k_s^S k_{on,S}^D k_{off,S}^T}{k_h^S k_{off,S}^D k_{on,S}^T}\right) = \ln\left(\frac{K_e^S}{K_e} \frac{k_h^{*S} k_s^*}{k_s^{*S} k_h^*} \frac{\alpha}{\alpha_{eq}}\right) \quad \text{(S22)}$$

b) $TS \rightarrow T^*S \xrightarrow{\text{hydrolysis}} DS$, after measurement, i.e. with bound substrate $S$; $D \xrightarrow{\text{exchange}} T^* \rightarrow T$, after resetting, i.e. without bound substrate. In analogy to point a):

$$\frac{\Delta S}{k_B T} = \ln\left(\frac{k_+^S k_h^{*S} k_{ex}^{D\to T^*} k_-}{k_-^S k_s^{*S} k_{ex}^{T^*\to D} k_+}\right) = \ln\left(\frac{K_e^S}{K_e} \frac{k_h^{*S}}{k_s^{*S}} \frac{\alpha}{\alpha_{eq}} \frac{[in]_{eq}}{[out]_{eq}} \frac{K_e k_s^S k_{on,S}^D k_{off,S}^T}{k_h^S k_{off,S}^D k_{on,S}^T}\right) = \ln\left(\frac{K_e^S}{K_e} \frac{k_h^{*S} k_s^*}{k_s^{*S} k_h^*} \frac{\alpha}{\alpha_{eq}}\right) \quad \text{(S23)}$$

c) $TS \xrightarrow{\text{hydrolysis}} DS$, after measurement, i.e. with bound substrate $S$; $D \xrightarrow{\text{exchange}} T$, after resetting, i.e. without bound substrate. Also in this case, we get the same $\Delta S$.

d) $TS \xrightarrow{\text{hydrolysis}} DS$, after measurement, i.e. with bound substrate $S$; $D \xrightarrow{\text{exchange}} T^* \rightarrow T$, after resetting, i.e. without bound substrate. Again, $\Delta S$ has the same value also for this set of operations.
